# Multiple groups of methanotrophic bacteria mediate methane oxidation in anoxic lake sediments

**DOI:** 10.1101/2021.11.24.469942

**Authors:** Guangyi Su, Jakob Zopfi, Helge Niemann, Moritz F. Lehmann

## Abstract

Freshwater lakes represent an important source of the potent greenhouse gas methane (CH_4_) to the atmosphere. Methane emissions are regulated to large parts by aerobic (MOx) and anaerobic (AOM) oxidation of methane that are important sinks in lakes. In contrast to marine benthic environments, our knowledge about the modes of AOM and the related methanotrophic microorganisms in anoxic lake sediments is still rudimentary. Here we demonstrate the occurrence of AOM in the anoxic sediments of Lake Sempach (Switzerland), with maximum in situ AOM rates observed within the surface sediment layers in presence of multiple groups of methanotrophic bacteria and various oxidants known to support AOM. However, substrate-amended incubations (with NO_2_^−^, NO_3_^−^, SO_4_^2–^, Fe^3+^ and Mn^4+^) revealed that none of the electron acceptors previously reported to support AOM enhanced methane turnover in Lake Sempach sediments under anoxic conditions. In contrast, the addition of oxygen to the anoxic sediments resulted in an approximately tenfold increase in methane oxidation relative to the anoxic incubations. Phylogenetic and isotopic evidence indicate that both Type I and Type II aerobic methanotrophs were growing on methane under both oxic and anoxic conditions, although methane assimilation rates were an order of magnitude higher under oxic conditions. While the anaerobic electron acceptor responsible for AOM could not be identified, these findings expand our understanding of the metabolic versatility of canonically aerobic methanotrophs under anoxic conditions, with important implications for future investigations to identify methane oxidation processes. Bacterial AOM by facultative aerobic methane oxidizers might be of much larger environmental significance in reducing methane emissions than previously thought.

## Introduction

Methane (CH_4_) is a powerful greenhouse gas in the atmosphere and a major portion of this gas in aquatic and terrestrial ecosystems is produced biologically in anoxic environments (e.g., anoxic waters and sediments) by anaerobic methanogens. In contrast to oceans, freshwater habitats such as lakes cover only a small portion of the Earth’s surface (0.9%; Downing et al. 2006), yet, they contribute a significant part to the global emissions of methane to the atmosphere (6-16%; Bastviken et al. 2004). The comparatively low methane concentrations in both surface and bottom waters of lakes (Blees et al. 2015; Donis et al. 2017) relative to those in anoxic sediments (mM range, e.g. Blees et al. 2014) suggest that methane in lake sediments is largely consumed before released to the water column. However, the primary sink for biogenic methane in anoxic lake sediments is still not well understood, and the physiological mechanisms and modes of benthic methane oxidation by microbes have not yet been unraveled.

In freshwater lakes, high activities of methane oxidation is often thought to occur at the oxic-anoxic transition zones (Lidstrom and Somers 1984; Kuivila et al. 1988; Frenzel et al. 1990; Bender and Conrad 1994; He et al. 2012). Here, methane oxidation is usually carried out by aerobic methane-oxidizing bacteria (MOB), and the methane oxidation with oxygen as electron acceptor (MOx) is considered the prime methane sink in lacustrine environments, and thus mitigator of lacustrine methane emissions. In anoxic lake waters, this pathway can be fueled by oxygen diffusion or intrusion events from the oxic water column (Hanson and Hanson 1996; Bastviken et al. 2002; Pasche et al. 2011; He et al. 2012; Oswald et al. 2016), or through oxygenic photosynthesis in shallow lakes (Oswald et al. 2015; Milucka et al. 2015; Rissanen et al. 2018). True anaerobic oxidation of methane (AOM) in anoxic environments is usually performed by anaerobic methanotrophic archaea (ANMEs) that are able to couple methane oxidation to the reduction of sulfate (Boetius et al. 2000; Orphan et al. 2002; Knittel et al. 2005; Milucka et al. 2012), nitrate (Haroon et al. 2013), metal oxides (Beal et al. 2009; Ettwig et al. 2016; Cai et al. 2018; Leu et al. 2020) or in a few cases, organic electron acceptors such as humic substances (Scheller et al. 2016). Methane oxidizing microorganisms may be metabolically quite adaptable. For example, recently identified anaerobic methanotrophic archaea in lake sediments, which are phylogenetically closely related to *Candidatus* Methanoperedens nitroreducens (i.e., nitrate-dependent methanotrophs or ANME-2d, Haroon et al. 2013), were reported to also perform methane oxidation coupled to sulfate reduction (Su et al. 2020). In addition to archaeal AOM, AOM can be performed by bacteria. Bacteria of the NC10 phylum (i.e., *Candidatus* Methylomirabilis), for example, can couple aerobic methane oxidation to nitrite reduction (Raghoebarsing et al. 2006; Ettwig et al. 2008), whereby the bacteria produce oxygen for methane oxidation intracellularly via the dismutation of nitric oxide (Ettwig et al. 2010). Thus, nitrite-dependent AOM can employ a pathway similar to that of canonical aerobic methane oxidation, which involves soluble and/or particulate methane monooxygenase enzymes. Such denitrifying methanotrophic bacteria have been recently reported for both lake sediments (Kojima et al. 2012; Deutzmann et al. 2014) and water column (Graf et al. 2018; Mayr et al. 2020). Irrespective of the microbial players involved, in anoxic lake sediments, pathways of AOM like nitrate/nitrite-dependent methane oxidation may be masked by aerobic process, due to the close proximity of the nitrate/nitrite- and oxygen-consumption zones near sediment-water interfaces, thus making it difficult to distinguish between true AOM and aerobic methane oxidation.

Methane oxidation may be ubiquitous in strictly anoxic lake sediments (Martinez-Cruz et al. 2018), yet, evidence for the occurrence of true AOM (i.e., by the above-mention AOM-performing microbes, involving electron acceptors other than oxygen) is still sparse (Schubert et al. 2011; CROWE et al. 2011; Sivan et al. 2011; Norði et al. 2013; Weber et al. 2017; Bar-Or et al. 2017; Su et al. 2020). Moreover, the role of traditionally aerobic methanotrophs in anoxic sediments and their contribution to AOM is still not well understood. Under oxygen limitation, some gamma-proteobacterial MOB (i.e., *Methylomonas* and *Methylomicrobium*) seem capable of oxidizing methane with nitrate/nitrite as terminal electron acceptors (Kits et al. 2015a; b), however, the initial attack of methane by methane monooxygenase still seems to be dependent on oxygen. Similarly, *Methylobacter* was recently shown to utilize methane for lipid synthesis during long-term anoxic incubations although the responsible electron acceptors for AOM were not investigated (Martinez-Cruz et al. 2017). High abundances of *Methylobacter* were also observed in the deep anoxic hypolimnion of a permanently stratified lake, in association with high methane oxidation rates (Blees et al. 2014), yet, the exact mechanism or mode of this apparent AOM by aerobic methane oxidizers, and the electron acceptor involved remained enigmatic. Finally, another putatively gammaproteobacterial lineage, *Crenothrix* was shown to have the potential to catalyze methane oxidation coupled the reduction of nitrate to N_2_O under oxygen-deficient conditions (Oswald et al. 2017). It appears that aerobic methanotrophs are versatile in their electron acceptor requirements, capable to conduct anaerobic respiration in oxygen-deficient or anoxic environments.

In this study, we investigated methane oxidation in anoxic sediments of Lake Sempach in Switzerland, with the particular goal to elucidate the potential electron acceptors and microbial players involved, and to verify whether the apparent anoxic benthic methane sink represents true AOM. Towards this goal, we combined sediment pore water hydrochemical data with in situ AOM rate measurements using radio-labeled methane, as well as slurry incubation experiments to study the impact of alternative oxidants (i.e., sulfate, iron, manganese, nitrite and nitrate) on AOM. In addition, we characterized the microorganisms that are involved in methane oxidation using approaches including lipid biomarker and 16S rRNA gene sequencing.

## Materials and Methods

### Study Site and sampling

Lake Sempach is a eutrophic lake located in central Switzerland. For more than three decades, the hypolimnion of this lake has been artificially aerated to maintain bottom water conditions oxic during summer thermal stratification, and to support mixing throughout the water column in winter (Stadelmann 1988). Alternative oxidants for methane oxidation (e.g. nitrate, nitrite, Fe^3+^, Mn^4+^) are found in the shallow sediments and porewaters. In March 2015, sediment cores (inner diameter 62 mm) were recovered with a gravity corer from the deepest site of Lake Sempach. Samples for dissolved methane concentrations were collected onsite with cut-off syringes through pre-drilled, tape-covered holes in the sediment core tube. 3 mL of sediment samples were fixed with 7.0 mL 10% NaOH in 20 mL glass vials, which were then immediately sealed with thick butyl rubber stoppers (Niemann et al. 2015). Porewater was extracted by centrifuging the 1 or 2 cm segments of a second sediment core under N_2_ atmosphere, and filtering the supernatant through 0.45 µm filters. Sample aliquots for sulfide concentration measurements were fixed with Zn acetate (5%) immediately after filtration. Sample aliquots for dissolved iron (Fe^2+^) and manganese (Mn^2+^) were fixed with HCl (6 M). The fixed or untreated filtered sample aliquots were then stored at 4 °C. Sediment samples for DNA extraction and particulate iron/manganese analyses were collected from a third sediment core, also at 1- or 2-cm intervals, and stored frozen at −20 °C until further processing.

### Chemical analyses

Methane concentrations in the headspace of NaOH-fixed samples were measured using a gas chromatography (GC, Agilent 6890N) with a flame ionization detector, and helium as a carrier gas. Nitrite was determined colorimetrically using the Griess reaction (Hansen and Koroleff (1999). Concentrations of sulfate, nitrate and ammonium were analyzed by ion chromatography (881 IC compact plus, Metrohm, Switzerland). Samples for dissolved iron (Fe^2+^) and manganese (Mn^2+^) were analyzed using inductively coupled plasma optical emission spectrometry (ICP-OES). Sulfide concentrations were analyzed in the laboratory photometrically (Cline 1969). Reactive iron oxide in the solid phase was extracted using 0.5 M HCl and then reduced to Fe(II) with 1.5 M hydroxylamine. Concentrations of Fe(II) were then determined photometrically using the ferrozine assay (Stookey 1970). Particulate reactive iron was calculated from the difference between the total Fe(II) concentrations after reduction, and the dissolved Fe(II) in the filtered sample. A total carbon analyzer (Shimadzu, Corp., Kyoto, Japan) was used to quantify dissolved inorganic carbon (DIC) concentrations. The penetration depth of oxygen into the surface sediments was determined using a Clark-type microelectrode sensor (Unisense A/S, Denmark).

### AOM rate measurements

To determine depth-specific AOM rates in Lake Sempach sediments, the subcore incubation approach was applied (Su et al. 2019). Briefly, subcores were collected by inserting three small plexiglass tubes (inner diameter 16 mm) into the fresh sediment core, and both ends of the core tubes were sealed securely with black rubber stoppers. After ∼1 h preincubation, 20 µL of ^14^CH_4_ solution (∼2.5kBq) was injected through each of the predrilled, silicone-sealed ports at 1.5 cm intervals. Subcores were incubated at in situ temperature (4 °C) in the dark for 2 days. At the end of the incubation, subcores were extruded, and subsamples were collected at 1 or 2 cm segments, and directly transferred into 100 ml vials containing 20 ml of the aqueous NaOH (5% w:w) to terminate microbial activity. Radioactivities of different ^14^C pools in the incubation samples and AOM rates were determined according to Su et al. (2019).

### Substrate-amended slurry incubations

#### Experiment 1 (sulfate, iron and manganese-addition experiments with ^14^C-labelled methane)

Anoxic slurries of ∼8 L were prepared by purging a mixture of sediments (surface 10 cm × 4 sediment cores, ∼1500 cm^3^) and autoclaved anoxic artificial sulfate-free medium (Su et al. 2020). Homogenized slurries of ∼100 mL were subsequently dispensed into 120-mL serum bottles and sealed with butyl rubber stoppers. For the electron acceptor amended incubations, stock solutions of sulfate, ferrihydrite and birnessite were added to final concentrations of 2 mM, 5 mM and 5 mM, respectively. For the inhibitor-addition experiment, molybdate was added to a final concentration of 4 mM. Base-killed controls (pH > 13) were also prepared and incubated in parallel. All bottles were further purged with N_2_ to remove any traces of oxygen. After 24-h preincubation, 1.5 ml pure CH_4_ was injected into the headspace of each bottle, which was gently shaken for 24 h, resulting in a final concentration of 100 µmol/L in the slurries. Finally, all incubation bottles were transported to an anoxic chamber and filled bubble-free with anoxic medium (100 µmol/L CH_4_) before injecting ^14^CH_4_ gas tracer. At different time points (30 and 60 days), incubation samples were stopped by adding 5 ml saturated NaOH after creating 20 ml headspace of N_2_. AOM rates were analyzed and calculated as described previously (Su et al. 2019).

#### Experiment 2 (nitrite-, nitrate-addition and oxic experiments with ^13^C-labelled methane)

To further test specifically the role of nitrite/nitrate in AOM, we performed slurry incubation experiments with a mixture of sediment and anoxic bottom lake water taken in November 2020. Briefly, 100 ml mixed slurries (1.5 g dw sediment per 120-mL bottle) were degassed with He to remove any traces of oxygen and background methane before transfer to an anoxic bag with continuous N_2_ flow. 5 mL of pure ^13^C-labelled methane (99.8 atom %, Campro Scientific) was injected into the headspace of the bottles, which were amended with anoxic stock solutions of nitrite and nitrate, with final concentrations of 0.48 mM and 2.4 mM, respectively. Incubations took place at room temperature under N_2_ atmosphere. Nitrite and nitrate were replenished, when consumed or when they remained at low concentrations. Killed controls (autoclaved and ^13^CH_4_), live controls (only ^13^CH_4_) and controls with only ^13^C-labelled bicarbonate were also incubated in parallel under the same conditions. For oxic experiments, slurries were spiked with pure oxygen and then injected with 5 mL of pure ^13^CH_4_. Oxygen was replenished regularly to maintain oxic condition in the incubation bottles. At different time points, incubation slurries were homogenized and 5 mL of the supernatant was collected under N_2_ atmosphere, and filtered with a 0.45 µm membrane filter for subsequent sulfate, nitrate, nitrite, DIC concentration and stable carbon isotope ratio analyses to determine the transfer of ^13^C into the product DIC pool. For the latter, 0.2-1 mL of water sample was transferred into a 12 mL He-purged exetainer (Labco Ltd) containing 200 µL zinc chloride (50 % w/v), and then acidified with ∼200 µL 80% H_3_PO_4_. The liberated CO_2_ was then analyzed in the headspace using a purge-and-trap purification system (Gas Bench) coupled to a GC-IRMS (Thermo Scientific, Delta V Advantage). Absolute ^13^C-DIC abundances were determined from the DIC concentrations and the ^13^C/^12^C ratios converted from δ^13^C-DIC values (Oswald et al. 2015). The temporal change in ^13^C-DIC with incubation time was then used to calculate slurry-incubation-based potential methane oxidation rates. Temporal solute-concentration and 13C-DIC changes were monitored over a total experimental period of 48 days. Thereafter, incubations were continued without further subsampling. After a total incubation period of approximately 5 months, slurry incubations were sacrificed for lipid analysis and DNA extraction (see below).

### Microbial lipid extraction and analysis

At the end of the incubation with ^13^CH_4_ (Experiment 2), after 160 days, triplicate slurries were combined and freeze-dried. Samples were then homogenized for lipid extraction using an Accelerated Solvent Extraction system (ASE 350, Dionex Corp., Sunnyvale, CA, USA). Samples were extracted with a solvent mixture of dichloromethane and methanol (9:1, vol/vol) over three extraction cycles at 100 °C and maximum pressure of 1500 psi. Total lipid extracts (TLEs) were collected in 60 mL glass vials, and concentrated using a vacuum evaporator system (Rocket Synergy Evaporator, Genevac Ltd., Ipswich, UK). Prior to extraction, an internal standard mix (5α-Cholestane, Nonadecanoic acid and 1-Nonadecanol) was added to each sample for the quantification of individual biomarkers. The TLEs were further evaporated to dryness, and then saponified with methanolic KOH-solution (12%) at 80 °C for 3 h. Neutral compounds were extracted by liquid-liquid extraction using hexane. After extracting the neutral fraction, fatty acids (FAs) were extracted from the remaining aqueous phase with hexane after acidification (pH ∼ 1). The fatty acid fraction was methylated using BF_3_ in methanol (14% v/v, Sigma Aldrich) at 80 °C for 2 h, and analyzed later as fatty acid methyl esters (FAMEs). Neutral compounds were further separated into hydrocarbon and alcohol fractions over silica glass gel columns, as described by Sicre et al. (1994) with the following modifications. The neutral fraction was dissolved in heptane and transferred onto the wet column. Aliphatic, cyclic and aromatic hydrocarbons were eluted with 2 mL heptane and 2 mL heptane/toluene (3:1 v/v), ethers and ketones with 2 mL heptane/toluene (1:1 v/v) and 2 mL heptane/ethyl acetate (95:5 v/v), and finally alcohols with 2 mL heptane/ethyl acetate (85:15 v/v) and heptane/ethyl acetate (80:20 v/v). The alcohol fraction was methylated with 100 µL pyridine and 50 µL bis(trimethylsilyl)trifluoracetamide (BSTFA) at 70 °C for 1 h.

All fractions were quantified using a TRACE GC Ultra gas chromatograph equipped with a flame ionization detector (GC-FID) (Thermo Scientific, Waltham, MA, USA). Individual lipid compounds were quantified by normalization to the internal standard and identified by comparing their retention times to those of laboratory standard mixtures, or by a gas chromatograph mass spectrometry (GC-MS, Thermo Scientific DSQ II Dual Stage Quadrupole) and the acquired mass spectra were identified through comparison with published data. Compound-specific stable carbon isotope ratios of individual compounds were determined using a gas chromatograph with split/splitless injector, connected on-line via a GC-Isolink combustion interface to a ConFlo IV and Delta V Advantage isotope ratio mass spectrometer (GC-IRMS, Thermo Scientific, Bremen, Germany). Both concentrations and δ ^13^C values of lipids were corrected for the introduction of carbon atoms during derivatization.

### DNA extraction, PCR amplification, Illumina sequencing and data analysis

DNA was extracted from both samples of Lake Sempach core sediments, as well as from the slurry sediments at the end of incubations (Experiment 2), using a FastDNA SPIN Kit (MP Biomedicals) following the manufacturer’s instructions. A two-step PCR approach was applied in order to prepare the library for sequencing at the Genomics Facility Basel. Briefly, a first PCR (25 cycles) was performed using primers 515F-Y and 926R targeting the V4 and V5 regions of the 16S rRNA gene (Parada et al. 2016). Sample indices and Illumina adaptors were added in a second PCR of eight cycles. Purified indexed amplicons were finally pooled at equimolar concentration into one library, and sequenced on an Illumina MiSeq platform using the 2×300 bp paired-end protocol (V3 kit). After sequencing, quality of the raw reads was checked using FastQC (v 0.11.8) (Andrews 2010). Paired-end read merger (usearch v11.0.667_i86linux64) was used to merge forward and reverse reads into amplicons of about 419 bp length, allowing a minimum overlap of 40 nucleotides and a mismatch density of 0.25. Quality filtering (min Q20, no Ns allowed) was carried out using PRINSEQ (Schmieder and Edwards 2011). Amplicon sequence variants were determined by denoising using the UNOISE algorithm (UNOISE3, usearch v11.0.667_i86linux64) and are herein referred to as ZOTU (zero-radius OTU). Taxonomic assignment was done using SINTAX v11.0.240_i86linux64 (Edgar 2016) and the SILVA 16S rRNA reference database v128 (Quast et al. 2013). Subsequent data analyses were carried out with Phyloseq (McMurdie and Holmes 2013) in the R environment (R Core Team, 2014) (http://www.r-project.org/).

### Data availability

Raw reads are deposited at the NCBI Sequence Read Archive (SRA) in the BioProject PRJNA759259, and can be accessed under the accession numbers SRR15689022-SRR15689034.

## Results

### Rates of AOM and geochemistry of Lake Sempach sediments

Concentration of methane in the sediments increased with depth, reaching a maximum of ∼1.75 mM at 10 cm depth. Methane oxidation was observed throughout the sediment core. Activity decreased with depth, with the maximum AOM rates (68.0 ± 19.0 nmol cm^-3^ d^-1^) measured in the upper 0-2 cm (Fig. 1B). Nitrate was only detected above 3 cm (up to 5 µM) and the sulfate concentration decreased downward from 78 µM at the sediment-water interface to 7 µM at 10 cm. Both sulfide and nitrite were below the detection limit at all depths. Reducible iron and manganese oxides were most abundant in surface sediments (24.3 and 30.8 µmol g^-1^ sediment respectively), and concentrations of both dissolved iron (Fe^2+^) and manganese (Mn^2+^) increased steadily with depth. Even under the condition of fully oxygenated bottom waters, oxygen was rapidly consumed in the uppermost sediment layers and penetrated to a maximum depth of 3 mm (Fig. 1F).

**Figure 1.**
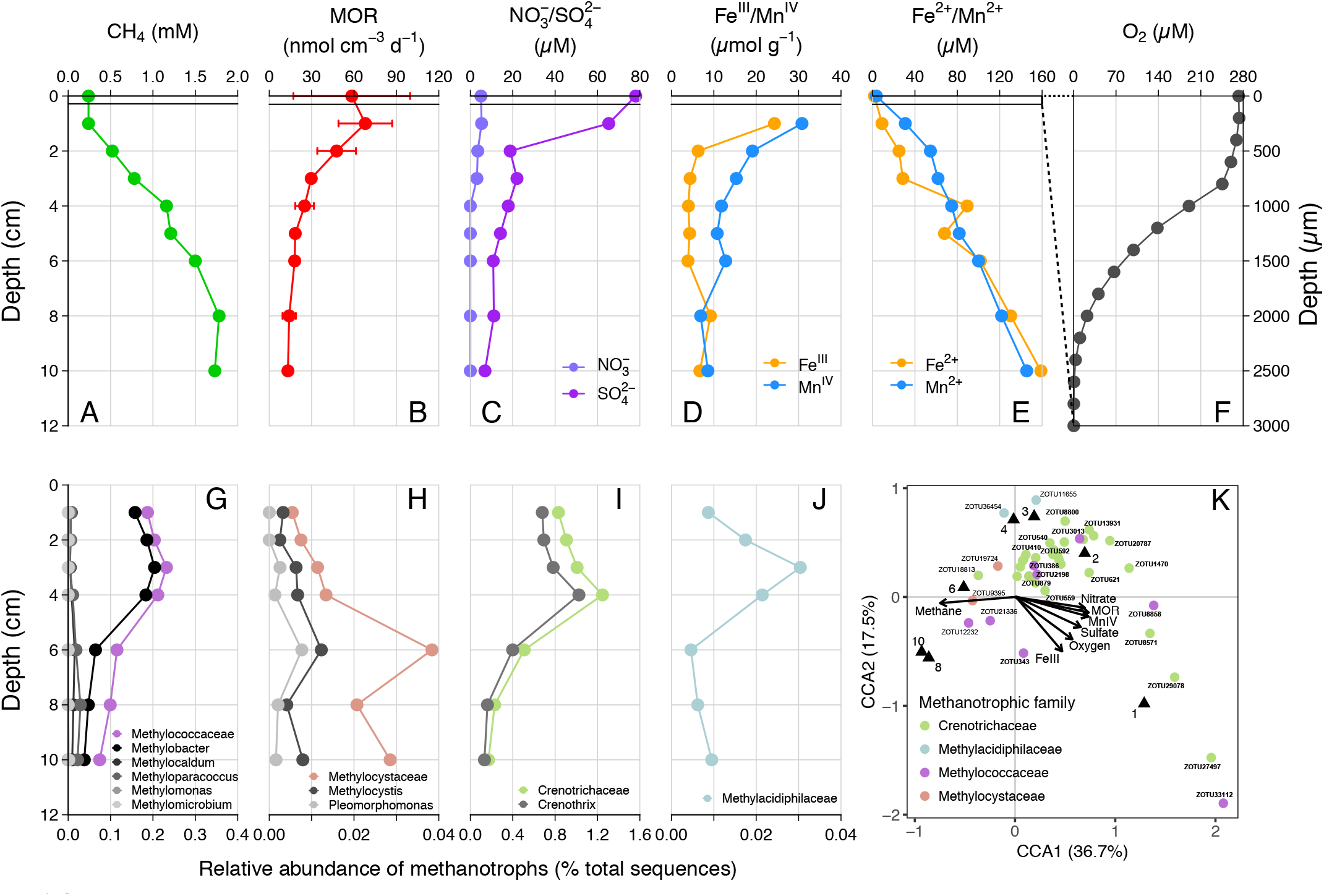
(A-F) Depth profiles of methane, methane oxidation rates, porewater nitrate/sulfate, particular iron(III)/manganese(IV) oxides, dissolved iron(II)/manganese(II) and oxygen in the sediment of Lake Sempach. Error bars represent standard deviations of triplicate rate measurements. No nitrite was detected. (G-J) Relative abundances of known families OF metahnotrophes and their subgroups at the genus level: (G) Methylococcaceae or Type I gamma-proteobacterial MOB, (H) Methylocystaceae or Type II alpha-proteobacterial MOB, (I) Crenotrichaceae, (J) Methylacidiphilaceae, (K) Canonical correspondence analysis (CCA), showing the relationship between MOR and methanotrophic species and the effect of nutrient concentrations on their distribution. ZOTU abundances were normalized and Hellinger-transformed prior to CCA analysis. A total of 42 ZOTUs accounting for > 92% of the methanotrophic community were shown on the plot and bold text represents taxa belonging to the genus *Methylobacter* or *Crenothrix*. Black triangles denote sediment samples at different depths.

### Diversity and abundance of methanotrophs in lake sediments

Analysis of sediment microbial community, based on 16SrRNA gene sequencing, revealed the presence of multiple groups of aerobic methanotrophs, including Type I gamma-and Type II alpha-proteobacterial MOB (Fig. 1G-H). Type I gamma-MOB consist of *Methylococcaceae*, of which the genus *Methylobacter* was the dominating cluster. The relative abundances of *Methylococcaceae* range from 0.07% to 0.23%, with peak abundance occurring at 3 cm depth. In comparison, Type II alpha-MOB *Methylocystaceae* was found at much lower relative abundances, with a maximum of 0.04% observed at 6 cm depth. A large number of 16S rRNA gene sequences (0.17– 1.25%, Fig. 1I), belonged to *Crenotrichaceae*, of which *Crenothrix* represented the predominant genus and the amplified sequence variants shared very high sequence similarities (96-100%) with what has been described recently as major methane oxidizers in stratified lakes (Oswald et al. 2017). In addition, a very small proportion of sequences (up to 0.03%) were affiliated with *Methylacidiphilaceae* within the *Verrucomicrobia* phylum. *Methylacidiphilaceae* were reported to perform aerobic methane oxidation in acidic geothermal environments (Islam et al. 2008; Carere et al. 2017). A canonical correspondence analysis (CCA) was performed to show any given correlation between the concentrations of potential electron acceptors, methane oxidation rates and specific methanotrophic species (Fig. 1K). The CCA triplot shows that members belonging to *Crenothrix* or *Methylobacter* were positively correlated to the methane oxidation rates, and negatively correlated to methane concentrations. Interestingly, the plot also demonstrates that *Crenothrix* and *Methylobacter* were more likely to be found in surface sediments with higher concentrations of potential electron acceptors (i.e., oxygen, nitrate, sulfate, iron and manganese).

### Effect of potential electron acceptor addition on AOM rates

To investigate the potential electron acceptors for AOM, sediments of the top 10 cm (i.e., where in situ rates were measured; Fig. 1B) were incubated with different electron acceptors. In the first experiment with ^14^C-labelled methane, no apparent increase in AOM rate was observed with any potential electron acceptor amendment, compared with the live controls (no electron acceptors added, Fig. 2A). In fact, incubations of both 30 and 60 days revealed very similar potential AOM rates between the untreated controls and amendments with sulfate and molybdate. Interestingly, the addition of iron oxides to the slurries did not stimulate methane oxidation but rather resulted in systematically lower AOM rates relative to the control experiments. Even more surprisingly, in incubations with added manganese oxide, methane oxidation was completely inhibited and substantial amounts of sulfate was produced. In the second experiment using ^13^C-labelled methane, we further tested the role of nitrate/nitrite in AOM, in direct comparison with aerobic methane oxidation after O_2_ injection. Similar to ^14^C-labelled methane incubation experiments, AOM occurred in the anoxic live controls (without any electron acceptors added). Incubations amended with nitrate and nitrite showed similar trends with respect to the control, hence, there was no obvious stimulation of methane oxidation by either of the NOx compounds (Fig. 3). For both untreated controls and nitrate/nitrite amendments, AOM rates were low during the first four days of the incubation period, but then increased dramatically between 4 and 8 days. Thereafter, they remained relatively low again (i.e., only subtle ^13^DIC accumulation between Day 8 and the end of the incubation). In contrast, methane oxidation was linear throughout the incubation period under oxic conditions, and the rates were almost an order of magnitude higher than the maximum rate in the control and NOx-amended incubations. Strikingly, sulfate was fully depleted after 8 days in the live controls (but not in the killed control), whereas there was extensive sulfate production in incubation bottles amended with both nitrate or oxygen. When nitrite was added, net sulfate production was also observed, at least during phases (e.g., initially and between Day 17 and 40; Fig. 3D), and at lower rates.

**Figure 2.**
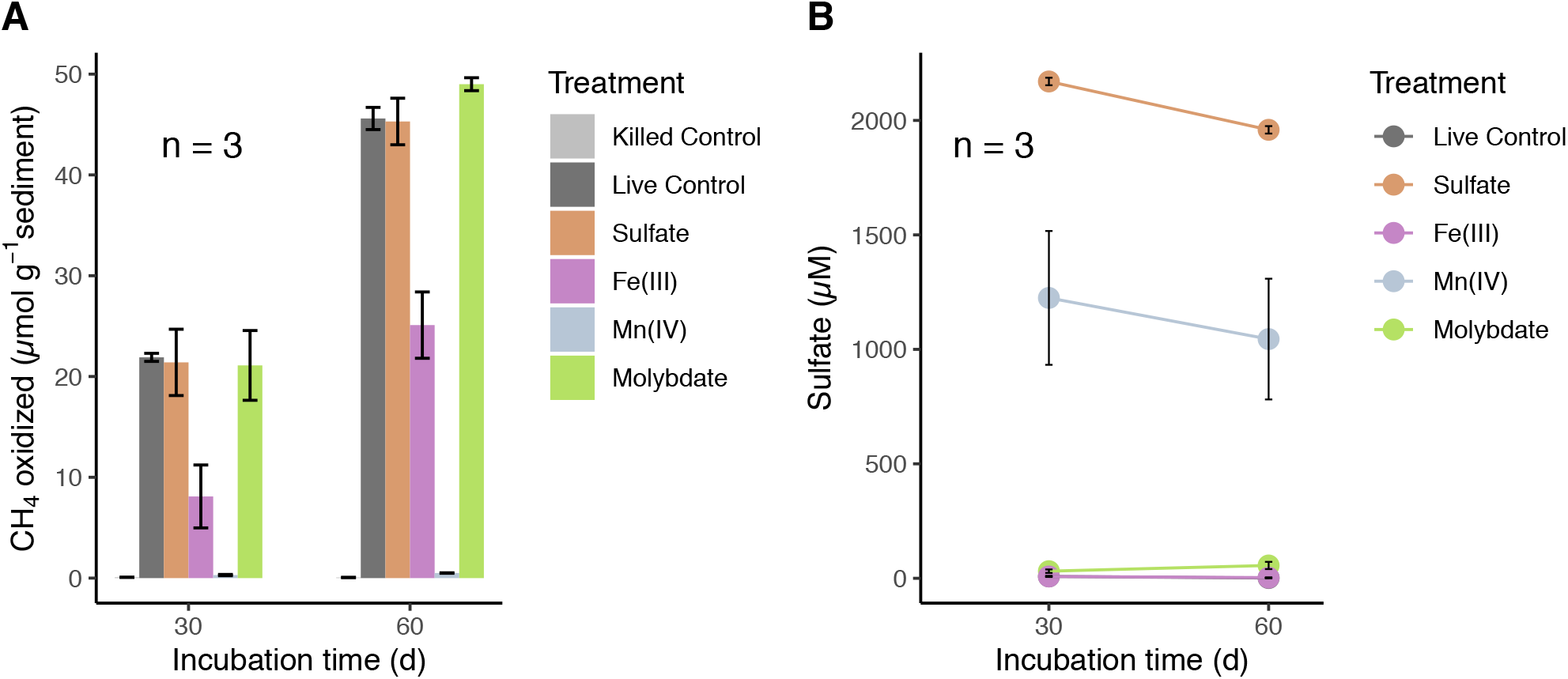
(A) Methane oxidation in anoxic slurries with ^14^C-labeled methane and different electron acceptors: Sulfate (2 mM), amorphous iron(III) oxides (5 mM) and manganese(IV) oxides (5 mM) after 30 and 60 days, respectively. (B) Sulfate concentrations in the incubation supernatants of different treatments after 30 and 60 days. Control incubations were conducted without electron acceptors added. No methane oxidation rates were detected in base-killed controls (pH>13). Error bars represent standard error of mean (n=3). Sulfate was below the detection limit in the live controls and in incubations with Fe(III) (points are overlapping).

**Figure 3.**
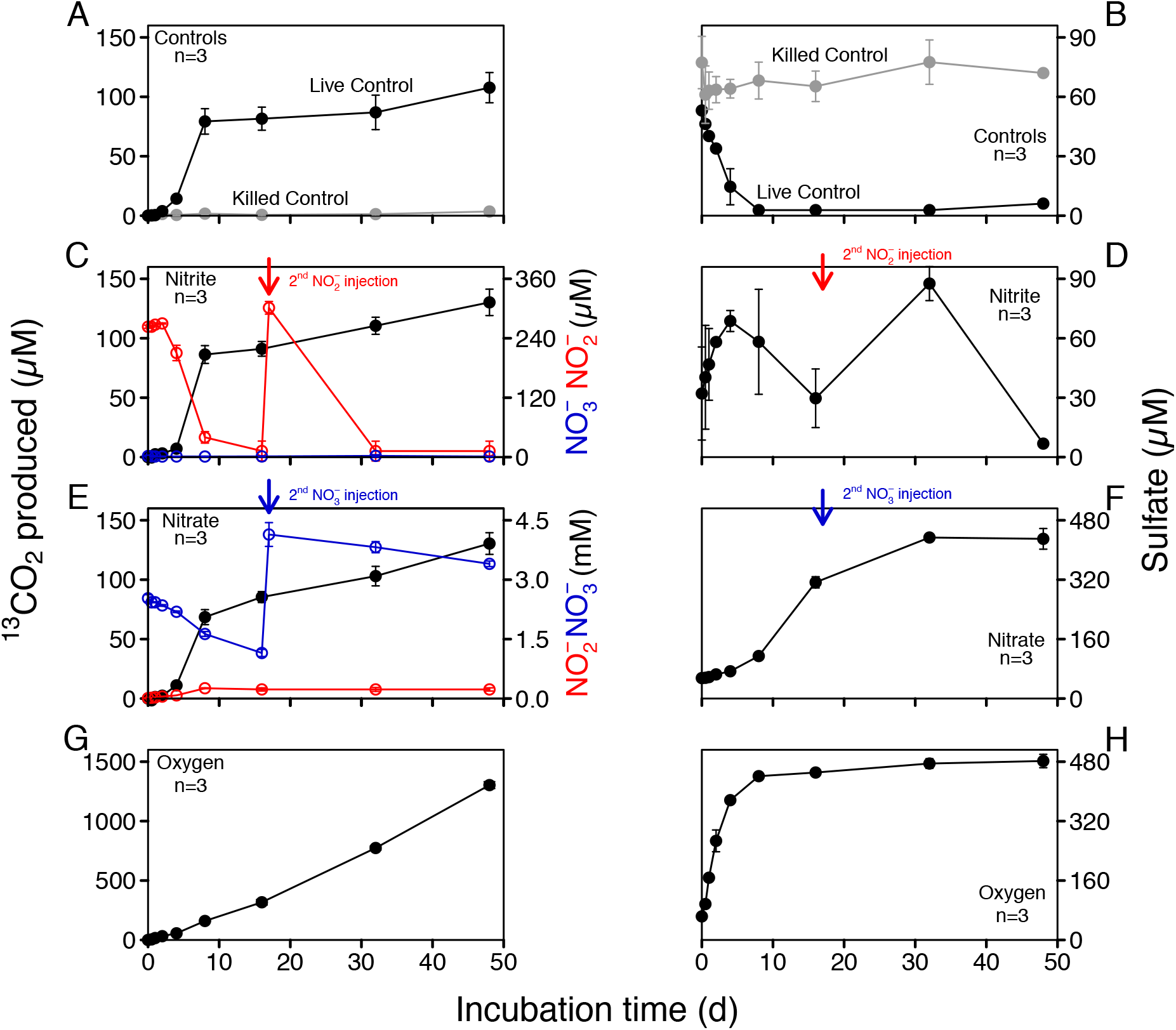
Concentrations of produced ^13^CO_2_ and sulfate in anoxic slurries amended with ^13^C-labeled methane and different electron acceptors (nitrite, nitrate and oxygen). (A, B) Controls including killed controls (autoclaved) and live controls (without electron acceptors added). (C, D) Nitrite addition; nitrite was replenished after 17 days. (E, F) Nitrate addition; nitrate was replenished after 17 days. (G, H) Oxic incubations. No nitrite and nitrate was detected in both controls and slurries with oxygen.

### Lipid biomarkers and microbial communities in incubations with ^13^CH_4_

At the end of the ^13^CH_4_ incubations (after160 days), the abundance and carbon isotopic composition of MOx/AOM diagnostic biomarkers (specific fatty acids and alcohols) were examined in the slurry sediments to constrain the methanotrophs involved and to infer carbon flow through the microbial community. The overall concentrations of lipid biomarkers in the different treatments were quite similar, except for incubations with oxygen, where concentrations were somewhat lower than in the other treatments (Fig. 4). In the untreated control and the nitrite-amended experiment, the most strongly ^13^C-enriched fatty acids were monounsaturated hexadecanoic acids (C_16:1ω7_ and C_16:1ω5_) and (ω-1)-OH-C_28:0_, with δ^13^C values ranging from 531‰ to 1558‰. By comparison, the highest ^13^C incorporation in incubations with nitrate was observed for (ω-1)-OH-C_28:0_ (1524‰), followed by 10-Methyl-hexadecanoic acid (10-Me-C_16:0_, 367‰). Most strikingly, δ^13^C analyses of biomarker extracts from the oxic incubations revealed significantly stronger ^13^C labeling of most fatty acids (compound-specific δ^13^C values between 6800 and 14500‰; Fig. 4) compared to the controls and to incubations with nitrite/nitrate. In order to assess concurrent autotrophic carbon assimilation, we performed additional slurry incubations with ^13^C-labelled bicarbonate only. No ^13^C incorporation at all was observed for (ω-1)-OH-C_28:0_ (−39‰), and the δ^13^C value of 10-Me-C_16:0_ (164‰) suggests only minor ^13^C-label incorporation. Generally, δ^13^C values of compounds diagnostic for methanotrophic microorganisms in incubations with ^13^C-bicarbonate were significantly lower than those in ^13^CH_4_ incubations with nitrite/nitrate (Fig. 4).

**Figure 4.**
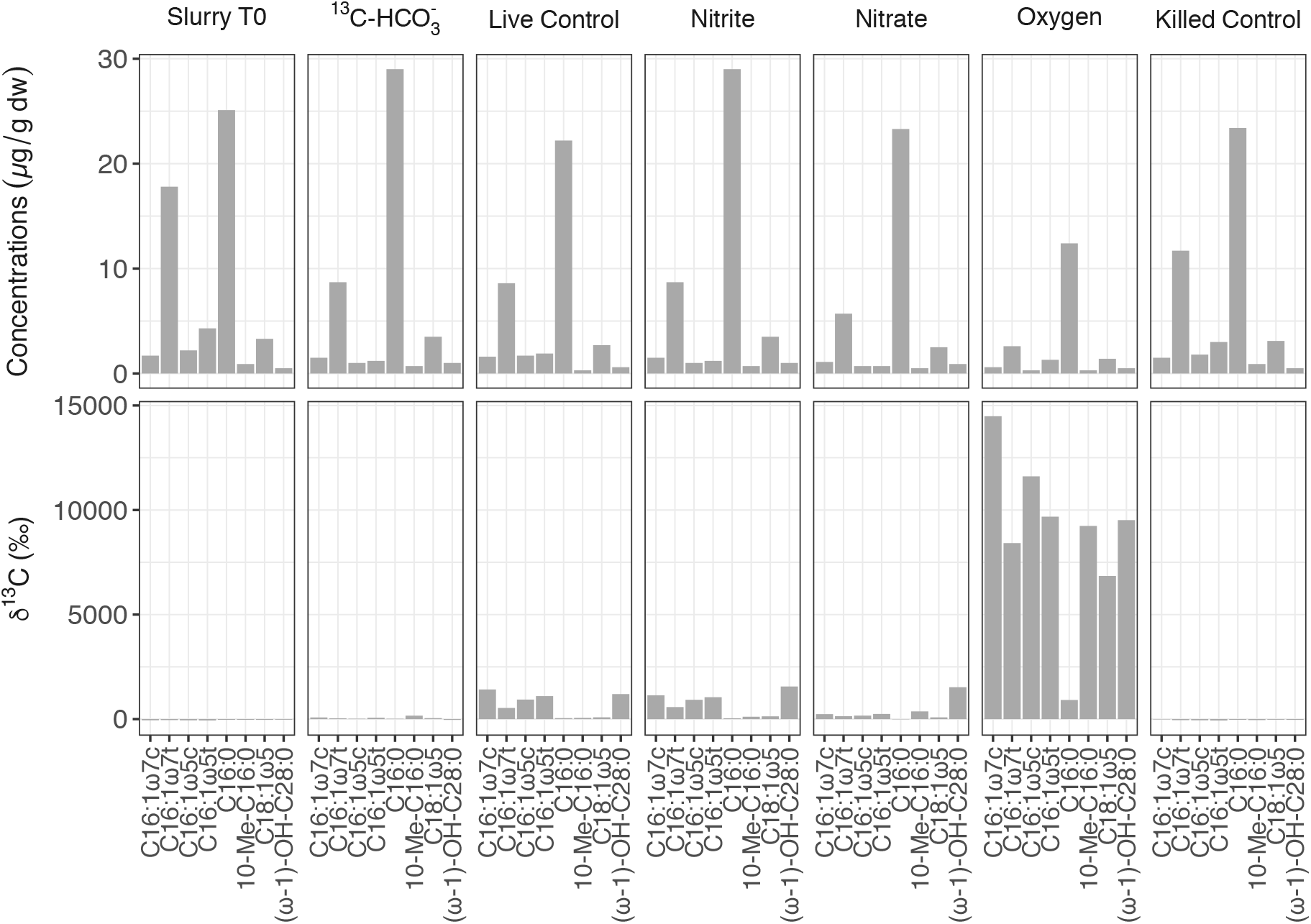
Concentrations and compound-specific δ^13^C values of fatty acids diagnostic for putative methanotrophs in the initial slurry sample (Slurry T0), and in ^13^CH_4_ incubations with different treatments after 160 days.

Initially (Slurry T0), *Euryarchaeota* was the most abundant archaeal phyla (Fig. 5A). The archaeal community structure in both the live control and incubation with ^13^C-bicarbonate, after 160 days of incubation, was similar to that of Slurry T0. However, No sequences of typical anaerobic archaeal methanotrophs such as ANME-1, −2, −3 and *Candidatus* Methanoperedens have been detected. As for bacterial composition at the class level (Fig. 5B), g*ammaproteobacteria* comprised a smaller proportion after nitrite/nitrate and oxygen additions. In oxic slurries, *Alphaproteobacteria* accounted for a larger proportion of the bacterial population compared to the pre-incubation microbial abundances. At family level, *Crenotrichaceae* and *Methylococcaceae* were the dominant groups among the known methanotrophs in the initial slurry sample (Fig. 5C), consistent with the diversity and abundance of methanotrophs in the sediment profiles (see Fig. 1G-J). After 160 days, *Crenotrichaceae* and *Methylococcaceae* remained dominant within the known methanotrophic families in all the treatments. Although representing a minor proportion, the relative abundances of *Methylocystaceae* slightly increased in incubations with nitrate/nitrite and oxygen when compared to Slurry T0. In oxic slurries, fractional abundance of *Methylococcaceae* was higher relative to Slurry T0 and the other treatments and the dominance of *Crenotrichaceae* seemed to be reduced (Fig. 5C). In addition, the anaerobic nitrite-dependent methanotroph *Candidatus* Methylomirabilis sp. was detected in incubation with nitrate and at a low abundance (1.36% of known methanotrophs; Fig. 5C).

**Figure 5.**
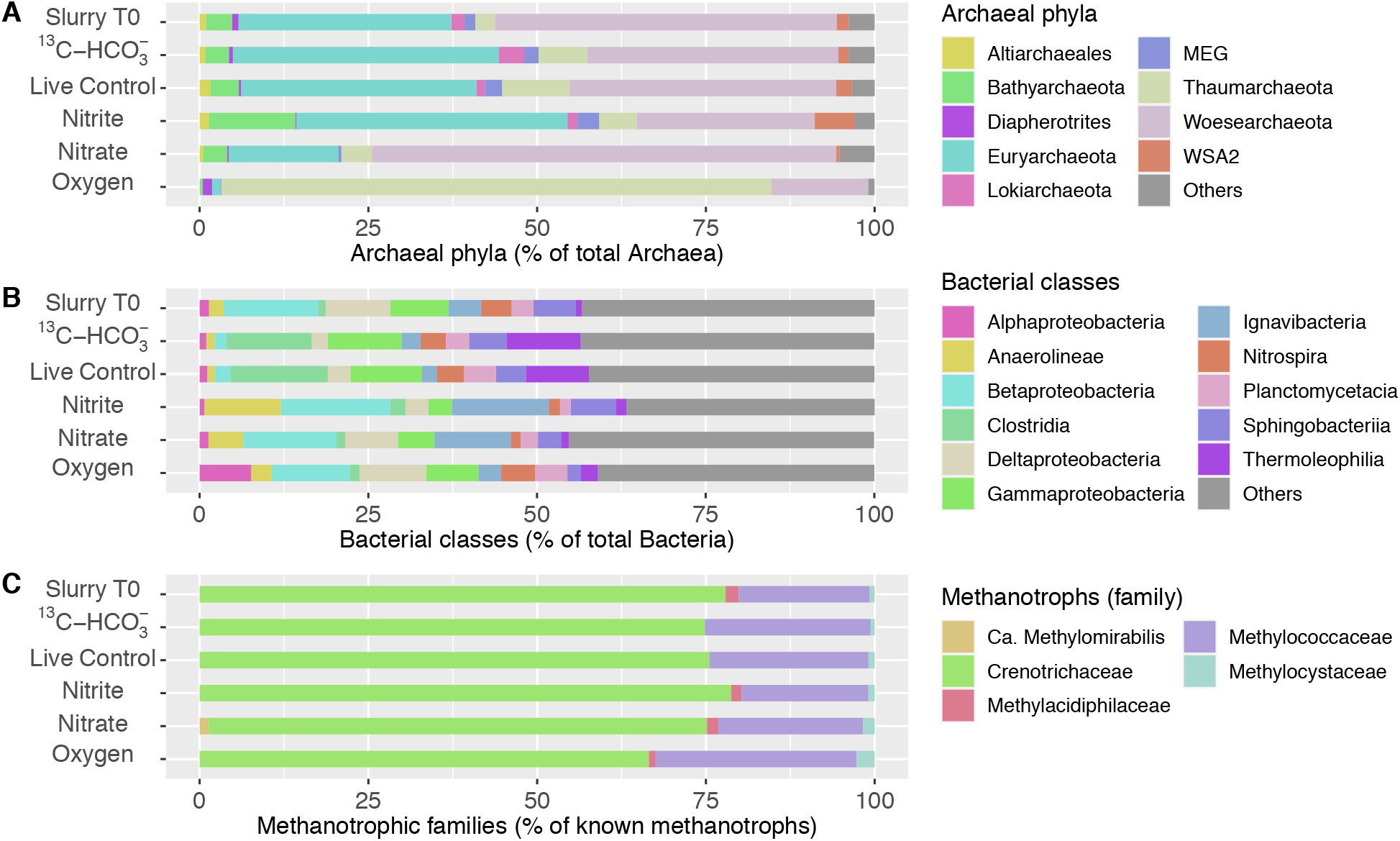
Microbial communities in slurry samples before (Slurry T0) and after incubation (160 days) with different treatments. (A) Relative abundances of the most abundant phyla within archaea. (B) Relative abundances of top bacterial classes. (C) Fractional abundances of known methanotrophic families. Data are based on read abundances of 16S rRNA gene sequences.

## Discussion

In Lake Sempach, despite the aeration of the deep hypolimnion, oxygen is rapidly depleted in the uppermost sediment layers (maximum O2 penetration depth of 3 mm). Based on the measured methane turnover rates as a function of depth (Fig. 1b), maximum methane oxidation activity was located at depths near the water-sediment interface, but apparent AOM (though at lower rates) was also indicated in the deeper, anoxic sediment layers. This is different from marine settings and some exceptional freshwater settings (e.g., Su et al. 2020) where highest AOM rates take place within the sulfate methane transition zones, well below the sediment surface (Iversen and Jørgensen 1985). Within oxic-anoxic transition zones of lake sediments, oxygen likely serves as the most important electron acceptors and aerobic methanotrophs play a dominant role in reducing the overall methane fluxes (Bender and Conrad 1994; He et al. 2012). In anoxic sediments, however, the pathway of methane oxidation with regard to the electron acceptors involved remains to be fully explored. Notably, the highest AOM rates in surface sediment layers corresponded to the concentration maxima of nitrate, sulfate, iron(III)-and manganese(IV) oxides, suggesting their potential roles for AOM. Despite the clear evidence of AOM in anoxic zones, no ANME-1 or 2, key microbial drivers of AOM in marine settings, were detected throughout the sediment core by 16S rRNA gene amplicon sequencing with the primers that match with a large fraction of the deposited sequences of anaerobic methanotrophic archaeal clades (Su et al. 2020). Neither could we detect molecular evidence for the presence of *Ca*. Methylomirabilis and *Ca*. Methanoperedens were detected either (Ettwig et al. 2010; Haroon et al. 2013; Su et al. 2020). Instead, the anoxic sediments contained high abundances of gamma-MOB and some alpha-MOB, which both are traditionally considered aerobic methanotrophs (Hanson and Hanson 1996). Most importantly, the sediments contained high relative abundances of *Crenothrix*, also gamma-MOB thought to be aerobes (i.e., requiring oxygen for methane activation), and regularly found as contaminant in drinking water systems (Stoecker et al. 2006). The relative abundances of *Methylococcaceae, Crenothrix* and *Methylacidiphilaceae*, respectively, were higher in upper sediment layers, while *Methylocystacea* were more abundant in lower portions of the sediments. The vertical distribution of various methanotrophic groups may indicate their different affinities towards methane, as evidenced, at least in part, by some studies showing that Type I methanotrophs were able to outcompete Type II MOB at low concentrations of methane (Amaral and Knowles 1995; Henckel et al. 2000). However, the presence of diverse aerobic methanotrophs in anoxic sediments led us to speculate that some of them, if not all, might have an alternative anaerobic lifestyle.

Indeed, increasing evidence has demonstrated the occurrences of gamma-MOB in anoxic lake waters and sediments (Blees et al. 2014; Milucka et al. 2015; Oswald et al. 2016; Martinez-Cruz et al. 2017). Also for Lake Sempach sediments, we were able to demonstrate active growth of gamma-MOB in absence of oxygen. That is, in O2-free incubations with Lake Sempach sediments and ^13^CH_4_,, the incorporation of the ^13^CH_4_-carbon into specific lipid biomarkers provides clear evidence for microbial methane assimilation. The monounsaturated fatty acids C_16:1ω7_ and C_16:1ω5_, which are typically indicative of Type I aerobic methanotrophs (Bowman et al. 1994; Hanson and Hanson 1996), were most enriched in ^13^C. In theory, the ^13^C enrichment could also be attributed to the assimilation of ^13^CO_2_ from methane oxidation into the biomass by autotrophic microorganism, as for example found for certain sulfate reducing bacteria (SRB) (Kellermann et al. 2012). Indeed, it has been reported that specific lipids such as C_16:1ω7_ and C_16:1ω5_ are associated with SRB that were involved in sulfate-dependent AOM at methane-rich seeps (Elvert et al. 2003). However, no or very low incorporation of ^13^C into these lipid biomarkers was observed in the incubation with ^13^C-labelled bicarbonate only. This indicates that the ^13^C-enriched lipids in the ^13^CH_4_ incubations were not derived from bicarbonate-assimilating SRB, but can rather be attributed to the direct assimilation of methane-derived carbon by Type I gamma-MOB. Direct uptake of ^13^C-labeled methane by alpha-MOB in the anoxic sediments was also observed, evidenced by the strong ^13^C-enrichment of long-chain (ω-1)-hydroxy fatty acids, such as (ω-1)-OH-C_28:0_. This lipid compound is mainly found in type II methane-utilizing bacteria including *Methylosinus* and *Methylocystis* (Skerratt et al. 1992). These are consistent with our findings regarding the composition of the methanotrophic community in the live control incubation slurry, which comprised relatively high abundances of both *Methylococcaceae* and *Methylocystaceae. Methylococcaceae* were dominated by *Methylobacter* and *Methylocystis* was the most abundant genus within *Methylocystaceae* (data not shown). However, we were not able to verify methane assimilation into the biomass of *Crenothrix*, because no diagnostic lipids have been reported for them to date.

Moreover, *Crenotrichaceae* likely also include bacteria that are not strict methanotrophs. Yet, due to the apparent capacity of at least some of the *Crenotrichaceae* genera to also oxidize methane anaerobically (Oswald et al. 2017), and given their high relative abundances in the anoxic sediments, however, we speculate that this methanotrophic group may comprise the most important methane consumers in Lake Sempach sediments. Overall, our molecular data and lipid biomarker analysis provide multiple lines of evidence that AOM in Lake Sempach sediments is most likely mediated by methanotrophs canonically considered aerobes.

According to the CCA triplot (Fig. 1K), members of *Crenothrix* and *Methylobacter* were more abundantly found at sediment with higher methane oxidation rates, and were associated with elevated concentrations of potential alternative electron acceptors in the anoxic sediments. However, in none of the electron acceptor-amended slurry incubations, i.e. neither with ^13^CH_4_ (nitrite/nitrate added) nor with ^14^CH_4_ (sulfate, iron or manganese added), stimulation of anaerobic methane turnover by any of the common anaerobic electron acceptors was indicated. This suggests an AOM pathway in Lake Sempach sediments that may be different from AOM reported for other investigated systems (Boetius et al. 2000; Orphan et al. 2002; Raghoebarsing et al. 2006; Vaksmaa et al. 2017; Su et al. 2020). In fact, the addition of metal oxides led to systematically lower AOM rates (iron) or complete inhibition (manganese). These observations appear, at first sight, at odds with recent studies showing the stimulation of both iron and manganese additions for aerobic methanotrophs under anoxic conditions (Oswald et al. 2016; Zheng et al. 2020). Yet, the inhibition of AOM as a result of excess “anaerobic electron acceptors” was also observed in previous studies (Segarra et al. 2013; Milucka et al. 2015; Rissanen et al. 2018). The addition of metal oxides to the slurries may change the redox condition of the systems, and may favor the growth of iron-or manganese-utilizing microorganisms that could outcompete methane oxidizers for essential nutrients, but further investigation is needed to reveal their effect on AOM. Notably, in ^13^CH_4_ incubations without electron acceptors added, the co-occurrence of increasing ^13^CO_2_ and decreasing sulfate concentrations over the first 8 days of incubation suggests a possible link between methane oxidation and sulfate reduction. On the other hand, the relatively large amounts of sulfate produced in nitrate-amended bottles, likely by the oxidation of sulfide with nitrate (Su et al. 2020), did not stimulate ^13^CO_2_ production significantly, when compared to the live controls, suggesting that sulfate was at least not a limiting electron acceptor for AOM in Lake Sempach sediments. The fact that sulfate reduction is not directly involved in driving AOM in Lake Sempach sediments is further supported by the observation that similar methane turnover rates were measured for incubations with and without molybdate, a competitive inhibitor for microbial sulfate reduction (Wilson and Bandurski 1958).

Despite the fact that aerobic methanotrophs are obligate aerobes, some of them seem to be able to survive long periods of oxygen starvation (Roslev and King 1995; Blees et al. 2014), and may even switch from respiring oxygen to nitrite or nitrate for methane oxidation (Kits et al. 2015a; b; Oswald et al. 2017). Both Type I MOB and *Crenothrix* were detected in our incubations, and their preference for O_2_ as oxidant of methane was clearly demonstrated for the Lake Sempach Sediments (Fig. 3G). However, how they (and/or other methane oxidizing microorganisms in the Lake Sempach sediments) drive methane oxidation in the absence of O_2_ remains uncertain. As outlined before, the AOM pathway involving nitrite or nitrate respiration seemed unlikely as neither enhanced methane oxidation rates significantly, indicating an as-yet-unidentified electron acceptor. While nitrite-dependent *Ca*. Methylomirabilis was not detected at all in any of the nitrite additions, it was present after incubation with nitrate addition, though at very low abundance. This finding is also consistent with the lipid data for the nitrate-amended anoxic incubations, which revealed a stronger 13C enrichment in the fatty acid 10-methylhexadecanoic acid (10-Me-C_16:0_), putatively diagnostic for *Ca*. Methylomirabilis oxyfera (Kool et al. 2012). It is plausible that these denitrifying methanatrophic bacteria contributed a small portion to the overall methane oxidation.

At this point, based on our slurry incubation results, we are not able to better constrain the electron acceptors involved in AOM in Lake Sempach sediments, and we can only speculate about what sustains the observed methane turnover in anoxic incubations. A recent study suggests that AOM can proceed in the absence of inorganic electron acceptors, via “reverse methanogenesis” (Blazewicz et al. 2012). This AOM pathway may offer a possible explanation for the observed “ineffectiveness” of all tested electron acceptors to stimulate AOM. Alternatively, other electron acceptors that were already present in the lake sediments (e.g., organic matter) could be involved. Indeed, humic substances have been recognized as a viable electron acceptor through the reduction of their quinone moieties (Lovley et al. 1996; Scott et al. 1999), and organic electron acceptors such as humic acids were suggested to play a role in AOM in wetlands (Smemo and Yavitt 2011; Blodau and Deppe 2012). In addition, anaerobic methanotrophic archaea in marine sediments were recently demonstrated to use humic-substance analogues (i.e., AQDS and humic acids) as electron acceptors in the absence of sulfate (Scheller et al. 2016). If humic (or similar organic) substances can indeed serve as methane oxidizing agent, and given the widespread distribution of these compounds in freshwater environments as important component of the sedimentary organic matter pools (Ishiwatari 1985), such mode of AOM may play an underappreciated role in reducing methane emissions anoxic lake sediments. Clearly, further investigations are required to explore their role in lacustrine AOM.

Most strikingly, additions of oxygen to our incubations with ^13^CH_4_ resulted in an approximately tenfold increase in ^13^CO_2_ production relative to the live controls or amendments with nitrite/nitrate. This clearly shows that aerobic methanotrophs were actively involved in methane oxidation when replenished with oxygen, and growth of these methanotrophic bacteria was evidenced by the substantial ^13^C-enrichment of diagnostic fatty acids. Our results thus highlight that there can be a strong potential for MOx in anoxic sediments, which, at least in Lake Sempach, seems to be mostly sustained by aerobic gamma proteobacteria (Type I MOB and *Crenothrix*). The fact that these microbes are found in sediments where O_2_ is fully depleted, and that they can resume full aerobic metabolic activity upon the addition of O_2_, attests to their resilience against O_2_ limitation. We propose that a potential capacity to switch to anaerobic modes of methane oxidation may be key to this resilience, allowing them to metabolize, at least at low level, also in the absence of O_2_.

Methane carbon assimilation rates in the oxic incubations were an order of magnitude higher than those in anoxic incubations. This is expected, since aerobic respiration provides much more energy for growth than respiration with alternative electron acceptors. While the result of the oxic incubation experiment confirm that aerobic methane oxidation is much more efficient than AOM in consuming methane, it needs to be noted that the vast majority of methane is produced, and accumulates, in the anoxic part of lake sediments. Here, AOM, just as in the marine environment, may serve as the important first step in a two-component benthic methane filter system. In this context of this ecosystem function, the metabolic versatility and apparent capacity of canonically aerobic methanotrophs to perform AOM has important implications for the natural mitigation of CH_4_ emission from lake waters to the atmosphere.

## Acknowledgements

We thank Thomas Kuhn and Judith Kobler Waldis for technical support in the lab. Thanks also go to Robert Lovas, Jana Tischer, Adeline Cojean and Yuki Weber for their great help during the campaigns on Lake Sempach. This research was supported by the China Scholarship Council (CSC) and additional funds from the Department of Environmental Sciences, University of Basel.

## Notes

### Competing Interest Statement

The authors have declared no competing interest.

